# HARLEY: A Semi-Automated detection of foci in fluorescence images of yeast

**DOI:** 10.1101/2021.11.29.470484

**Authors:** Ilya Shabanov, J. Ross Buchan

**Affiliations:** Department of Molecular and Cellular Biology, University of Arizona, Tucson, AZ 85721, USA

## Abstract

Quantification of cellular structures in fluorescence microscopy data is a key means of understanding cellular function. Unfortunately, numerous cellular structures present unique challenges in their ability to be unbiasedly and accurately detected and quantified. In our studies on stress granules in yeast, users displayed a striking variation of up to 3.7-fold in foci calls and were only able to replicate their results with 62-78% correlation, when requantifying the same images. To facilitate consistent results we developed HARLEY (Human Augmented Recognition of LLPS Ensembles in Yeast), a customizable software for detection and quantification of stress granules in *S.cerevisiae*. After a brief model training on ~20 cells the detection of foci is fully automated and based on closed loops in intensity contours, constrained only by the a-priori known size of the features of interest. Since no shape is implied, this method is not limited to round features, as is often the case with other algorithms. Candidate features are annotated with a set of geometrical and intensity-based properties to train a kernel Support Vector Machine to recognize features of interest. The trained classifier is then used to create consistent results across datasets. HARLEY is aimed at users without technical expertise, allows for batch processing and is freely available, which should be of broad interest to users focused on analysis of microscopy data in yeast.

## INTRODUCTION

Live and fixed cell fluorescence microscopy is a common approach utilized by biologists to elucidate understanding of cellular function. The ability to accurately identify and quantify objects (i.e., foci) in microscopy data is thus key, and yet routinely papers are published with microscopy data that suffers from a lack of rigorous, unbiased analysis of foci of interest. An accurate measure of foci number, size, intensity or geometry may often reveal variations in intriguing underlying biological phenomena. However, variation in the aforementioned foci properties, and in cellular background signal or other relevant cellular contexts, can make foci quantification resistant to efficient, automated, analysis. Several approaches to quantifying cellular microscopy data have thus been developed as briefly outlined below.

### Threshold based methods

Threshold based methods assume that features of interest have a higher or lower intensity than the background surrounding them. Identifying such thresholds leads to a binary segmentation in feature and background areas. Various morphological and denoising operations are usually combined to obtain a good segmentation.

Top-hat or H-maxima transformations are typical choices employed for denoising, detection or accentuation of features by algorithms like FoCo (Lapytsko et al., 2015), FociCounter (Jucha et al., 2010) or CellProfiler’s (McQuin et al., 2018) “Speckle Counting” pipeline. Statistical thresholding like Otsu’s method or many other alternatives are then used to yield a final threshold. These methods are readily extendable to 3D as demonstrated by FocAn (Memmel et al., 2020).

Morphological operations often require a kernel that presupposes a shape of the feature in question. In most cases a disc of a set radius is used, limiting the algorithm to detecting round foci. More importantly, thresholding-based techniques are sensitive to a non-uniformly distributed background signal. For example, in the analysis of stress granules (SGs) in *S.cerevisiae*, vacuoles, which have no cytoplasmic signal, and vary greatly in size, can impair the identification of the actual cytoplasmic background signal.

### Kernel based methods

In a very general sense kernel-based methods identify local features of an image by scanning through the image and scoring positions according to their similarity to an expected “kernel” (typically a small search pattern image). A convolution operation between kernel and image yields a similarity map. Portions of the image correlating most closely to the kernel receive high scores and thus can be identified as features of interest by using thresholding or local maxima detection.

FociQuant (Ledesma-Fernández & Thorpe, 2015) detects kinetochores (and other foci) and BUHO (Perez-Pepe et al., 2012) detects SGs using this general method.

The main limitations of kernel-based methods are the kernels themselves, since they offer a somewhat rigid representation of what a feature is, different shapes, rotations or scales require multiple kernels and may rely on weight or threshold parameters for the final result.

Convolutional Neuronal Networks (CNN) solve some of these problems by learning the shapes of the kernels from a dataset and performing convolutions at different scales. CNNs however require big amounts of labeled data to be trained and can be computationally expensive.

### Differential Scale-Space Methods

For a given image *I*(*x, y*) differential scale-space methods generally first construct a stack of images smoothened at different strengths (or scales) *t* called a scale space *L*(*x, y, t*) and then look for extrema of differential features (e.g., any combination of partial derivatives of *L*) therein. Detected maxima thus not only contain spatial coordinates of the feature, but also a scale corresponding to the size of the feature making them scale invariant.

The most common differential operators are the Laplacian of Gaussian (LoG), its approximation Difference of Gaussian and the Determinant of the Hessian Matrix (DetH). A significant number of other operators have been proposed described in (Lindeberg, 2012). The challenge with these methods arises in delimiting a boundary of the feature, since they inherently only detect points. One solution is to define a region that is convex around the detected point and define the boundary along the zero crossing of the Gaussian curvature as proposed by (Marsh et al., 2018).

These methods detect many false negatives that usually require a threshold parameter to be removed. Another problem with methods like LoG and DetH is that they yield the highest values for round features; irregular and elongated shapes therefore pose a problem in noisy data. Furthermore, in cell images with strong background signal the curvature of the cell outline itself starts contributing to a set of false negatives.

### Machine Learning Methods

Machine Learning approaches typically work on raw images and given labeled datasets learn automatically how to extract and classify the features of interest. As in our approach (described later) they can be easily combined with classical computer vision methods to increase performance or reduce the amount of data needed for training.

Many architectures have been proposed for various tasks, with CNNs being commonly used. DeepFoci (Vicar et al., 2020) utilizes U-Net, a specific CNN architecture to detect and segment foci and nuclei with promising accuracy in a fully unsupervised manner. A combined approach in (Hohmann et al., 2020) uses classical techniques to detect regions of interest, extracts an over abundant set of features using filters and feeds these results into a classifier. FindFoci (Herbert et al., 2014) uses a machine learning approach to devise parameters to filter out foci of interest in accordance with a human experimenter.

Deep Learning approaches can also leverage context. Having the whole image as input, information about location inside the cell and from foci to foci can be incorporated. Unsurprisingly, these approaches usually outperform previously mentioned methods. This however comes at a cost of long training times and the need for large, labeled datasets. To our knowledge there are no deep learning-based approaches specific to yeast SGs.

### Other methods

In theory many of the methods can be combined to create a more nuanced quantification. An example of such work is found with Obj.MPP software (Graeve et al., 2019) that uses a marked point process framework. A set of parametrically defined objects like disks, squares, or ellipses is compared via different quality functions to the image. These quality functions process differential (e.g., gradient) information along the edges of objects and around them. AutoFoci (Lengert et al., 2018) combines a top-hat transformation, an estimate of local curvature (using LoG) and a foci shape derived coefficient to create an Object Evaluation Parameter.

Several issues regarding analysis of microscopy data remain unresolved. First, excluding manual classification (e.g., in Fiji) or generic approaches (e.g., CellProfiler), the identification of SGs, or other cytoplasmic foci with significant cytoplasmic background signal has seen very few tailored automated approaches. Second, as reported in an exhaustive study on bioimaging informatics tools (Schneider et al., 2019), overall usability is a significant hurdle for the use of automated methods, making CellProfiler one of most popular, yet not optimal (Vicar et al., 2020) methods. Finally, in biological publications the issue of human variability and error is often not addressed. A previous study (Herbert et al., 2014) showed with three users that in manual counting of γH2AX foci, 20-30% percent of assignments are unmatched between any two experimenters. While there is no evaluation of how consistent the same human is and the use of the F1 score for unbalanced datasets can be somewhat misleading this study is very indicative of significant human bias. A similar study on γH2AX foci with 5 experimenters from 3 labs (Runge et al., 2012) reported 81.1% of assessments between experts to deviate in a statistically significant manner.

Here, we asked 7 users of varying degrees of self-identified expertise to quantify a yeast SG dataset twice, and observed a striking ~4-fold variation between users in terms of foci calls. Furthermore, users could only replicate ~60-80% of their individual quantification performance when re-quantifying a subset of the SG dataset. This motivated us to develop a new user friendly framework, cell and foci detection algorithm termed HARLEY (**H**uman **A**ugmented **R**ecognition of **L**LPS **E**nsembles in **Y**east) which enables manual or trained quantification of SG and other foci-like organelles. HARLEY is freely available on Github (https://github.com/lnilya/harley).

## MATERIALS AND METHODS

### Yeast SG induction and deconvolution microscopy

BY4741 yeast transformed with pRB1 (Pab1-GFP, Edc3-mCh) were grown to mid-log phase and subject to NaN3 stress as previous described (Buchan et al., 2011). Z-stack data (10 slices, 0.4μm) was collected using a 100X objective (NA 1.45) on a Deltavision Elite microscope and subject to deconvolution and maximum intensity Z-stack projection. A brightfield light image was also collected at the Z-stack center to facilitate cell boundary detection.

### Software versions

Our software is based on ReactJS 17 for its frontend and Python 3.8 for its backend. Exact versions of employed packages can be found in the respective package files in our Github repository.

### Recruitment of users and dataset

7 independent users with varying degrees of self-identified familiarity in quantifying SG data were recruited by the PI to quantify SGs in a collection of 55 individual cell images, using a web-based app we developed. (See the Labeling step of the Model Training pipeline in HARLEY; **Supplementary video 3**). This allowed users to both label and delimit a boundary around the periphery of each putative SG foci.

On a scale of 1:Layman, 2:Intermediate, 3:Expert the users only the user “R” self classified as expert, with all others being intermediates.

Foci candidates were determined using our contour loop method. After labeling a candidate (consisting of shortest and longest contour loop) as a focus users could choose a contour loop most closely matching their perception of foci outlines by dragging the mouse across the focus in an interactive UI. The same method is used in the model training step of our software and can be explored further there.

A second round of quantification of the same cells, randomized in order from the previous scoring, was used to assess intra-user quantification variability.

Two users (A and M) failed to delineate boundaries around foci and only clicked on the foci in question, we therefore excluded their results in foci size analysis.

### Model Evaluation using the Matthew Correlation Coefficient

For a set of identified foci candidates, a decision needs to be made whether a candidate is a feature of interest, leading to a binary classification problem with a heavily unbalanced dataset (i.e., several times fewer foci than non-foci). As (Chicco & Jurman, 2020) have described previously, the commonly employed measures of Recall, Precision or their combination the F1 score (also known as Dice coefficient) to compare these classifications are poorly suited when the dataset is unbalanced. To compare the results of models and users we therefore use the Matthews Correlation Coefficient (MCC) (Matthews, 1975), a much more robust measure for unbalanced datasets.

The MCC is defined as a function of the confusion matrix consisting of the four numbers:

- *TP* :True positives - number correctly classified foci
- *TN* :True negatives - number of correctly rejected foci
- *FP*: False positives - number of wrongly classified foci
- *FN*:False negatives - number of wrongly rejected foci

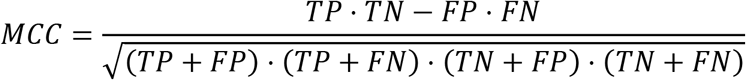

With the values ranging from −1 to 1. 1 being perfect classification, 0 being the correlation of two completely randomized experiments (e.g., coin toss) and −1 being perfect misclassification.

### Cross Validation MCC

When comparing the results of a user’s classification with the results of a model trained on this same data (i.e., the user’s model) it would be unwise to simply use the MCC, since an overfitted model would yield over optimistic scores.

We therefore use the mean of a 6-fold cross validation error over 6 models trained on the respective subsets of data. k-fold cross validation is a technique that splits the dataset into k disjunct sets of equal length *D* = ∪_*i*_ *D_i_* with *D_i_* ∩ *D_j_* = Ø ∀*i* ≠ *j*. We then train the model on the set *D_ni_* = ∪_*k*≠*k*_ *D_k_* and evaluate the MCC on the prediction of the remainder of the dataset *D_i_*.

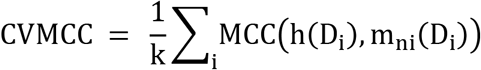

Denoting the user prediction and the model prediction trained on dataset *D_ni_* with *h*(*D_i_*) and *m_ni_*(*D_i_*) respectively.

The resulting correlation will be lower but a more generalized measure of model performance. We use this score only when comparing the user to their own model. In all other cases the model is trained on the whole data set and its prediction then compared using the regular MCC.

### Training of HyperParameters

Support Vector Machines use a regularization parameter C that governs the size of the margin separating the classes. Since we are using radial basis functions as our kernels, the parameter *γ* governs the size of this kernel.

A typical method to find these parameters is to optimize them with regards to the Cross Validation, since it avoids overfitting them to the data. We use a logarithmic grid search with values between 10^-3^ and 10^3^ for C and 10^-8^ and 10 for *γ* to find parameters that fit the data better. In practice we found this step to give only very minor improvements over the default settings of 1 and ‘scale’ for C and *γ* respectively from the sklearn.svm python implementation.

### Random Quantifier

We use the MCC with a random quantifier (Rnd) as a first measure of noise in a user’s quantification efforts. For each cell Rnd simply randomly picks the same number of foci as the user has chosen, resulting in a “classified” dataset which can now be compared to the user’s result using the MCC. This comparison is repeated 50 times and the mean is used as the final score, reported in this publication.

### Feature space

Our feature space is built by first obtaining the closed contours of a given length (as defined by experimenter and scale of images), then predicting the size of the foci with our algorithm and extracting a set of features based on expert knowledge from these values: 1.) maximum, minimum, mean intensity of focus; 2.) area, eccentricity and solidity; 3.) size in relation to cell size; 4.) brightness ratio predicted outline and center; 5.) brightness ratio longest outline and center; 6.) absolute difference in brightness center and outline; 7.) normalized distance to cell center; 8.) brightness index inside cell (1 brightest in cell, 0 dimmest in cell). These choices were also partially motivated by ease of extraction and speed of execution. The resulting system runs in real time along the user’s labelling giving instant feedback on the test and training errors and converges quickly in practical terms. The feature space is whitened and rescaled 0-1 before processing by the SVM.

### Extraction of cell boundaries from bright field images

Since cells in fluorescence microscopy may or may not have a varying degree of background signal that could be used to delimit single cells, we use brightfield images to extract cell outlines. The process yields a binary mask that is used with fluorescence images to extract and normalize data for single cells.

#### Ridge detection

Our algorithm first uses the Frangi ridge filter (Frangi et al., 1998) that is then thresholded to give the first draft of cell outlines. A graphical UI guides the experimenter throughout the whole process allowing easy correction of mistakes and cell-identifying parameters.

#### Cleaning

A cleaning step is then employed that removes foci candidates that do not match certain criteria. These criteria are size or area constraints, as well as bounds on eccentricity and solidity of blocks available in the skimage.measure v0.19.0 package. In practice, these parameters once set can be reused with very little modification from image to image.

#### Ellipse Fits

The cleaned outlines are then thinned (Zhang & Suen, 1984) to give a skeleton image of the outlines. We now evaluate a set of pixel positions *C* = {*x_i,j_*} in the image whose distance to the skeleton lies between *r*_min_ and *r*_max_; these are parameters on minimum and maximum cell size.

For each of these candidate points we fit an ellipse that approximates skeleton points around *x_i,j_* and calculate an approximation error as a sum of squared distances between the skeleton and the resulting ellipse. Additionally, we can filter out points where the skeleton lacks points in a range of angular directions, e.g., the outline is not a closed shape.

The result is a heatmap with local minima representing locations of cell centers. It is important to note that parameters defined here depend on the magnification of the images and are transferrable between images.

#### Boundaries

The final ellipses at these minima of approximation error are then “snapped” to the boundaries using a version of the active contours model (Kass et al., 1988) to yield final outlines. These are prone to noise and strong internal edges, like vacuoles or nuclei, resulting in only partial recognition of the cell. These errors can be manually rejected, by an interactive UI.

Even though this method works well **(Sup. Video 1)** and proposes a semi-automated way of detecting cells that in our test images is more than sufficient to gather and review hundreds of cells in minutes, we acknowledge that it probably will not outperform state of the art deep learning techniques. However, the typical number of cells quantified in liquid-liquid phase separation (LLPS) research is in the magnitude of hundreds and we therefore did not investigate better methods further.

### Preprocessing of Fluorescence Images

Fluorescence images are first stacked using a max-projection and then denoised using Non-Local Means Denoising (Buades et al., 2011). Both steps are guided through an interactive UI **(Figure 2B and Sup. Video 2)**. Since noise levels and image stacks vary between experiments and microscopes, these settings are selected image by image.

## RESULTS

### Manual quantification of yeast SG data generates large intra and inter-variability amongst users

7 users of intermediate to expert familiarity with SG quantification identified SG foci and their boundaries in a dataset of 55 yeast cells exposed to NaN3 stress. These users repeat their quantification 1-2 weeks later with a shuffled version of the dataset. Many users exhibited significant differences in foci identification **(Figure 1A)**. User foci sections varied from 21% to 79% (3.76-fold) of available foci (see Materials and Methods) on average **(Figure 1B)**, while average SG foci size selections ranged from 108 to 152 nm **(Figure 1C)**. Foci calls from the first and second quantification run by users resulted in a reproducibility range of 62-78% **(Figure 1D**; MCC, see Materials and Methods). These results demonstrate how volatile quantification between users can be, and that even the most conservative users (those who score fewer foci on average) have difficulty in reproducing their own quantification results. Finally, for most users we find only small variation in the determination of foci size between quantification runs **(Figure 1E)**.

**Figure 1:**
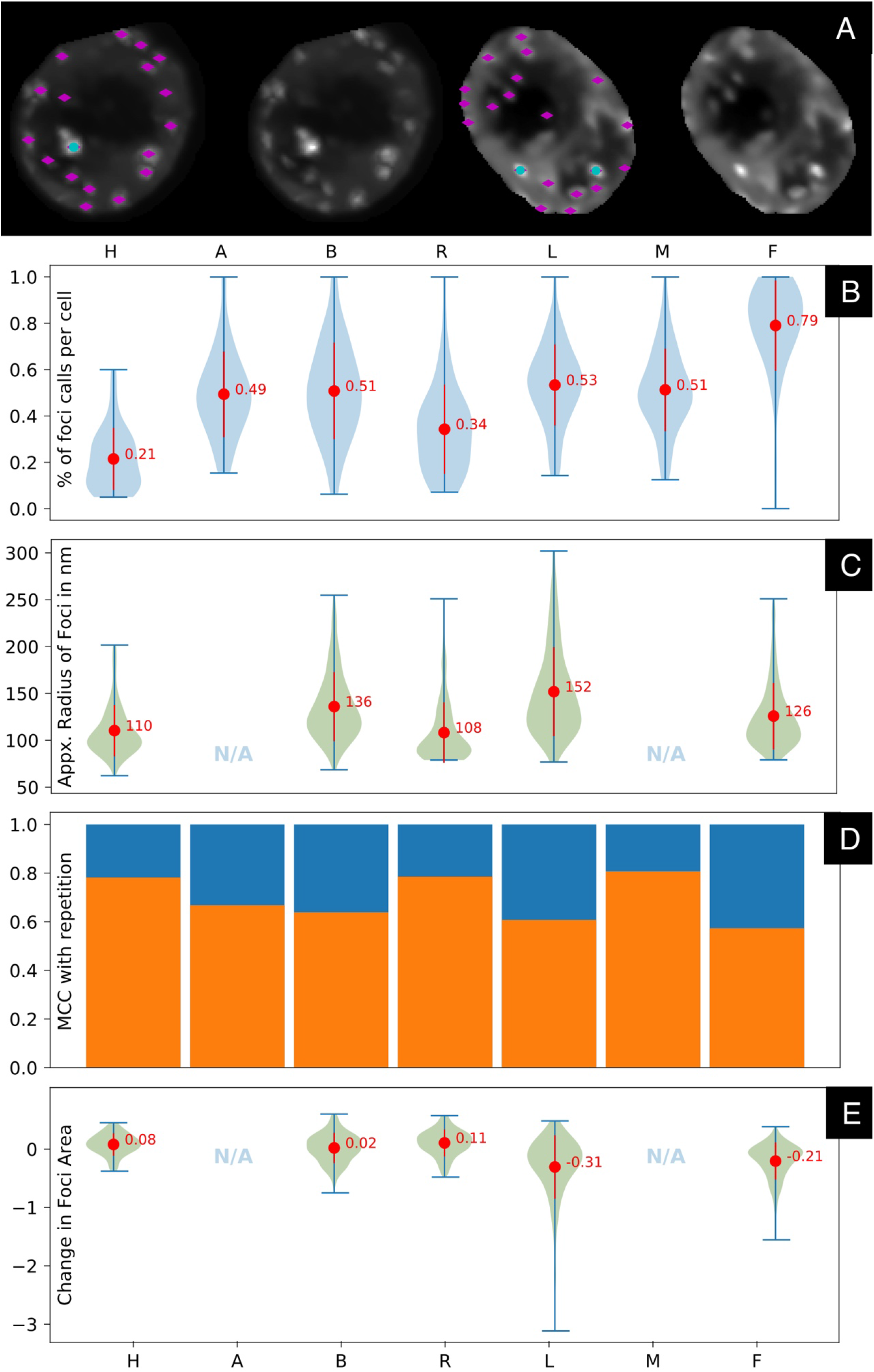
SG quantification is highly variable amongst users and on repetition. Seven users (H,A,B,R,L,M and F) classified foci. Users that did not specify foci sizes are excluded in panels C and E. **A:** Example of two ambiguous cells in dataset, with choices of the most permissive (magenta) and most conservative user (teal). **B:** Ratio of selected foci to available foci in individual cells by user. **C:** Approximate radius in nm of foci by user. Since foci are not perfectly, round, the radius corresponds to a circle of the same area as selected foci. **D:** MCC between two quantification runs on the same dataset (see Materials and Methods) **E:** Ratio of foci area differences in second run relative to the first.

The message from these findings is clear: Manual quantification is prone to very large errors especially between different users, particularly when the signal to noise ratio of SG foci to diffuse cytoplasmic background is not especially high (a common issue in SG yeast data analysis). Due to a lack of accessible, automated data analysis tools suited for SG quantification in yeast, we set about developing an automated software solution.

### HARLEY (Human Augmented Recognition of LLPS Ensembles in Yeast); a novel software for cell segmentation and foci identification via a trained model

Our software allows researchers to efficiently proceed from image data to final quantification results and consists of a pipeline of steps that are independent and interchangeable with other methods. At important steps of the pipeline a human experimenter may use the UI to adjust results. The full pipeline consists of 1.) Extraction of single cell boundaries as masks on bright field images. 2.) Max-intensity stacking and denoising of fluorescence images (preprocessing). 3.) Extraction of contour loops in individual cells as foci candidates 4.) Manual labelling of valid foci by a human to train a model and 5). Automated classification of foci using the obtained model **(Figure 2)**.

**Figure 2:**
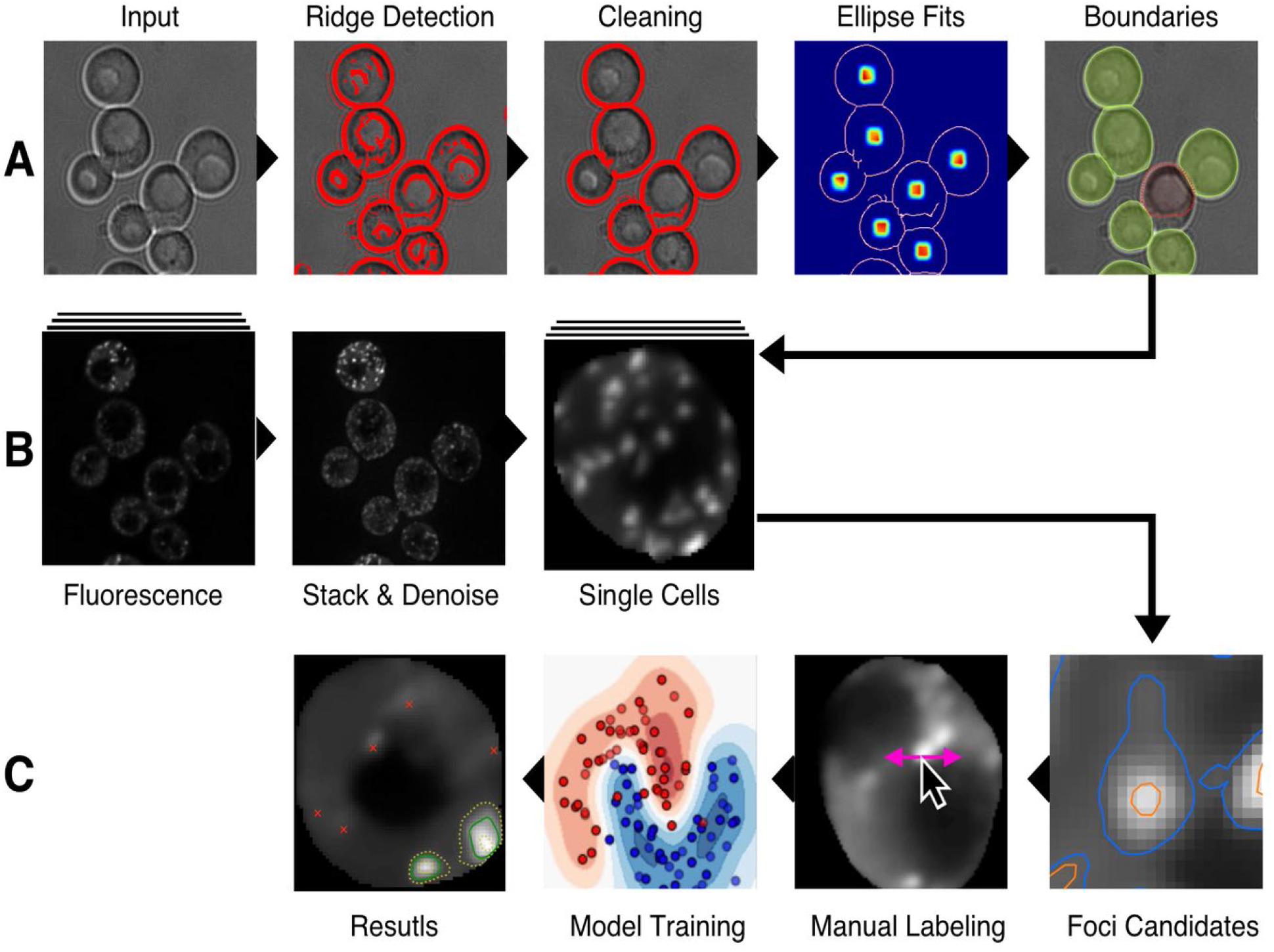
Schematic overview of HARLEY. **A:** Detection of cell outlines using brightfield images. **B:** Preprocessing and extraction of single cells using the cell outlines and stacked fluorescence images. **C:** Extraction of foci candidates as contour loops of constrained length, labeling of a training set and automated selection of foci via the trained model.

#### Identifying Foci Candidates and boundaries

The key building block of our algorithm are closed contour loops of length within a predefined range, readily found to sub-pixel precision using a marching cubes algorithm. If one were to imagine an image in 3 dimensions with the intensity being the height, closed contour lines are lines along which the intensity (or height) does not change and that either circle a peak or a valley, akin to topology lines on maps.

This analogy immediately allows us to identify an important property: Contour loops around a point never intersect. They are either non overlapping, equal or the longer one contains the shorter one, rendering it unnecessary to deal with clumped foci explicitly.

We can now define a focus-candidate as a tuple of an outer and a contained inner contour: (*C_k_, C_j_*) with the contour loops *k* ≠ *j* having the lengths *L*_max_ and *L*_min_ respectively; the lengths being parameters set by the experimenter **(Sup. Video 3)**. As we shall see later the algorithm is not very sensitive to these settings, they are simply upper and lower bounds on the features in question.

Each contour line has its respective intensity *I_k_* < *I_j_* and at this location an intensity *I_k_* < *i* < *I_j_* will describe a contour between the outer contour *C_k_* and the inner contour *C_j_*.

For close-by foci, an outer contour might contain multiple inner contours, in which case we treat those as multiple separate candidates. Later, when the final outline of foci are calculated, the two candidates will either merge into one or remain two separate foci. The problem of merged or clumped foci is therefore dealt with implicitly and does not require any explicit segmentation, like a watershed approach that is commonly employed in such scenarios (Hohmann 2020, McQuin et al., 2018).

#### Identifying the final contour from a focus candidate

Just as different users would draw contours of a focus differently (**Figure 1C**), there cannot be any “correct” method of solving the problem of finding an optimal contour for a candidate. We suggest that more importantly than whether the result looks correct to a given user, contour focus identification should consistently give the same results and not rely on parameters.

A good example of a parameterless approach is the Hessian Blob algorithm (Marsh et al., 2018) that uses the zero crossing of the gaussian curvature of an image as a boundary.

The Gaussian curvature of the image however requires a scale factor (of the Gaussian), which is a result of the blob detection algorithm but is not present in our approach. Discrete methods of curvature estimates can lack precision given the size of the features that is often only a few pixels across.

We defined our foci candidates as a tuple of two contours (*C_k_*, *C_j_*) an inner and an outer. The problem of finding final contour *C_opt_* between these two thus can be described as finding an intensity *I_k_* < *I_opt_* < *I_j_* that maximizes some function *f_k,j_* of the two contours: *I_opt_*(*k, j*) = argmax (*f_k,j_*(*I*))

The concept is visualized in **Figure 3C** with inner and outer contours displayed as yellow lines. The choice of the function to optimize is arguably ambiguous. However, in order not to introduce any parameters we examined differential properties. Inflection points and maxima are typical choices in the literature. We suggest using the intensity at which the curvature(i.e., the second derivative 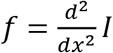) is at a maximum rather than the inflection point (where it has a zero crossing).

**Figure 3:**
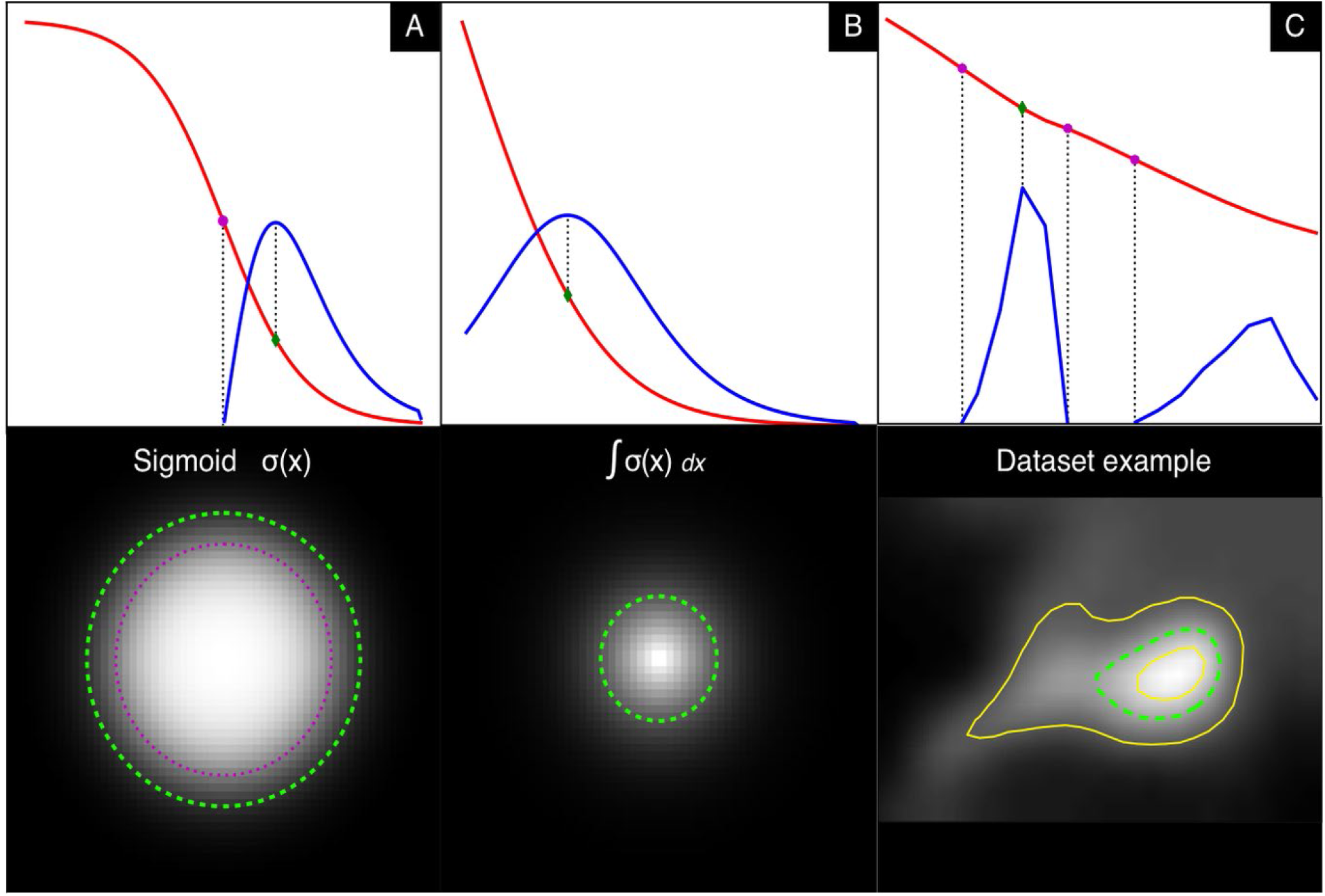
Intensities and curvatures (not to scale) of artificial and real foci with resulting images. **A and B:** Foci modelled by different functions in red and their second derivative in blue (not to scale). The top row represents a slice through the intensity along the radius and has the inflection point (magenta) and the curvature maximum (green) marked. These points correspond to outlines of foci depicted as circles of the same color in bottom row. **C:** Real life example from our dataset with multiple inflection points. Yellow lines represent inner and outer contours of the foci candidate C_k_ and C_j_ respectively.

To motivate this choice, we examine different models for an intensity peak by rotating a function of intensity around its y-axis **(Figure 3A-B)**. We see that it is easy to conceive a focus without any inflection points of intensity **(Figure 3B)**. Generally, the curvature maximum outline corresponds more closely to the human perception of foci, while the inflection point boundary appears somewhat too tight.

In the above examples the intensity is a function of radius *I*(*r*); what we have though are contour outlines along a fixed intensity that are not circular. In order to approximate a solution, we can imagine a circular focus where the intensity drops linearly with the radius and the area is therefore *a*(*I*) = *πr*^2^. We observe further that the area is now in a simple relation to the intensity function *I*(*r*) we are looking for:

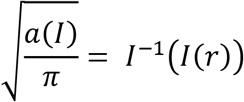

This means we can obtain the intensity function by inverting the square root of the area function of a contour, which leads to a correct solution in case of a circular focus, where the intensity equals the radius or an approximation otherwise. Evaluation of areas of contours between (*C_k_, C_j_*) at different intensities gives a piecewise linear function that can be inverted (since the area is a monotonous function) and differentiated using numerical methods to derive a final contour for this focus.

**Figure 3C** shows the result for a focus in our dataset. Note that in this case we have multiple inflection points, but only one absolute maximum, making the resulting contour intensity unambiguous. Furthermore, this example demonstrates the limited influence of parameters *L*_max_ and *L*_min_ for determining the final foci. While they need to be chosen reasonably to include those points of maximum curvature, they have no significant influence on the choice of the final focus outline making the algorithm more stable regarding these parameters.

#### Foci Classification

Having found foci as closed contour lines along the maximum curvature we end up with many more candidates, most of which are just perturbations of background signal rather than what human users would classify as *bona fide* foci. The choice of which candidates to count as foci and which to reject strongly depends on the user, as previously shown **(Figure 1B)**.

Typical criteria a user may intuitively use in identifying foci include a certain % of intensity above the surrounding background signal, foci morphology and co-localization or proximity with other identifiable cellular structures. Some of these criteria are hard to quantify and due to its subjective nature only of limited reproducibility. Defining these rules for all users and use cases is therefore not possible. Instead, we extract basic properties of the image and let a classifier algorithm deduce rules based on a labeling provided by a user.

To train an automated classifier we first extract a feature vector *x_i_* from each focus (see Materials and Methods) yielding a dataset *D* = {*x_i_* | *i* = 1,…, *N*}. Our software allows users to simply set bounds on these features to generate labels *l_i_* ∈ {0,1} where 1 and 0 mean that a closed contour loop is or isn’t a focus respectively.

More generally such rules are a plane that partitions the feature space:

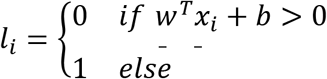

Given a manually labeled dataset we use a Support Vector Machine (SVM) to solve the problem of finding optimal weights *w* and *b* under the side constraint that the plane should be separating the two classes with as large a margin as possible. In simplified terms a kernel SVM extends this plane of separation to a non-linear surface of separation. In our approach we use radial basis functions as kernels (details on hyper parameter settings in the materials and methods). SVMs are a powerful and flexible class of supervised algorithms that have been successfully applied to image and microscopy data for decades (Miao 2016, Maglogiannis 2004, Wang 2017). We forego the formal mathematical explanation and implementation details. which are well described elsewhere (Chang & Lin, 2011).

In summary we have 1.) devised a method to detect potential foci using two contour loops of minimal and maximal length; 2.) proposed a method of finding an optimal contour within these two by approximating the intensity function from the area of contours; and 3.) proposed the use of a SVM on various features extracted from these foci to automatically and rapidly label valid foci. Our software deals with all the preparatory steps to this point, like cell outline detection, denoising and stacking of fluorescence microscopy images (**Figure 2, Supplementary videos 1-4**).

### Models match their human behaviour closer than other humans while generalizing towards a shared “ground truth”

While human users usually classify foci from a set of self chosen rules like “foci are 50% brighter than their surroundings” we have shown that they do not follow these consistently with the best correlations on reclassification of the same dataset being only around 80% **(Figure 1D)**. The result of user classification could therefore be thought of as a deterministic predictor *pr* plus some added random noise *ξ* that depends on the dataset and user.

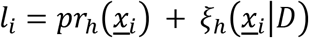

Intuitively a classifier capable of generalizing the data should primarily capture the deterministic term, leaving the noise term indicative of an upper bound for the model performance. Captured in the deterministic term is a user’s familiarity with what a SG should look like and that which a model should capture primarily. While our sample size is too small to undertake a thorough statistical examination of the noise and assess performance boundaries for the model, our results are indicative that our model indeed removes some of this noise.

For most users, the correlation to their model lies above the correlation to other users **(Figure 4A)**, which means that the model performs a quantification that closer matches that user versus another.

**Figure 4:**
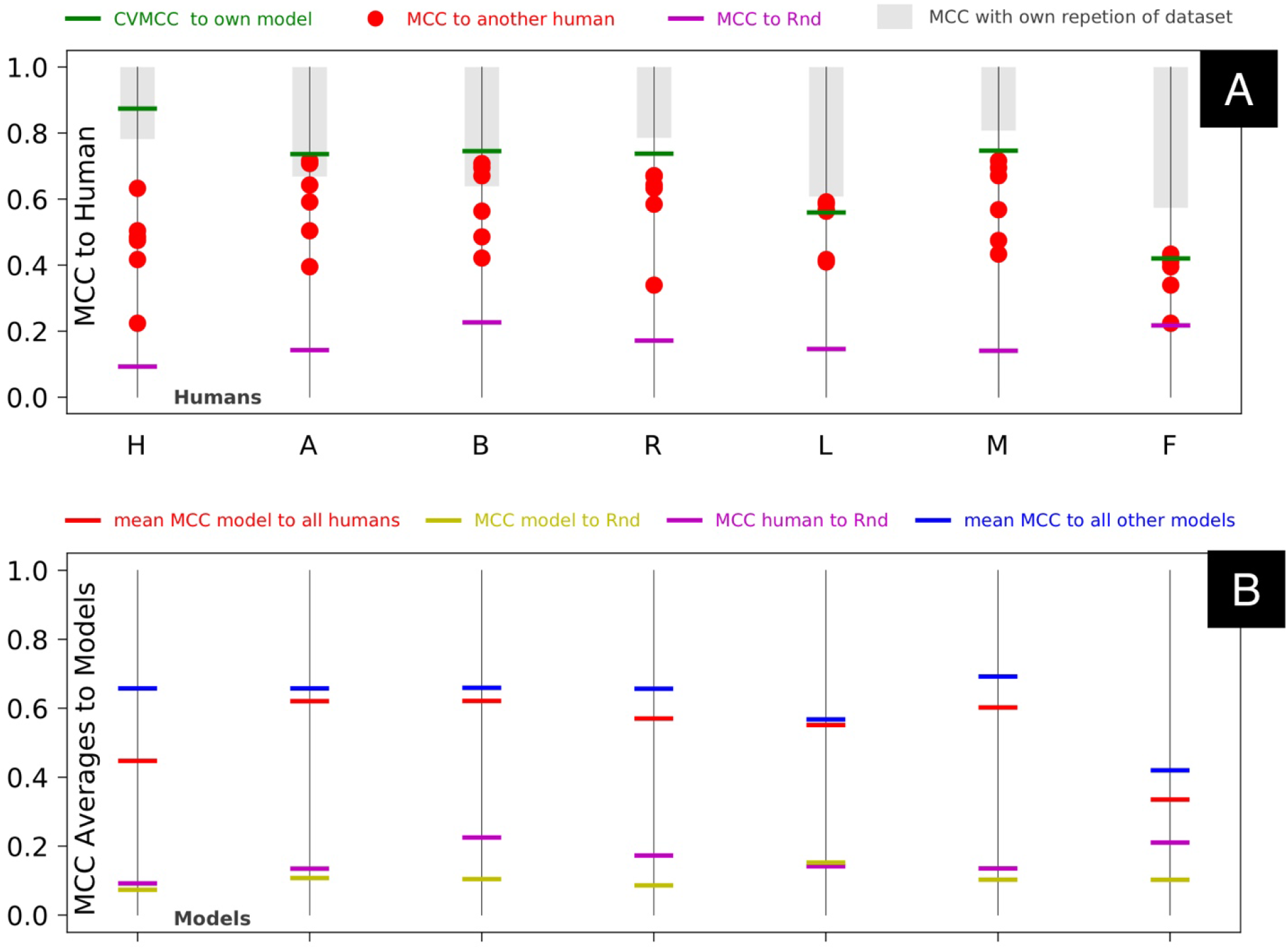
Evaluation of Model Performance. **A:** Correlation of users to the trained model (green), other users (red) and the random classifier (magenta). The gray bar goes from 1 to the correlation of the user with themself (see **Figure 1D**) **B:** Mean correlations of models to all other models (blue), all users (red) along with the correlations of this model with the random classifier (yellow) and for comparison the user’s correlation from panel A (magenta).

To assess the relation to noise, we compared the users and models to a random classifier (“Rnd”, see Materials and Methods). **Figure 4B** compares models trained on their respective user to other users and other models. Notably, models generally correlate closer together than to the other users (blue bars above red), regardless of the user they have been trained on. This suggests that models tend to generalise the deterministic term of the classification rather than learn the noise. In theory if the understanding of a SGs was identically defined across all users (like what a “Dog” or “Cat” is), then users would only differ in the noise term and all models would tend to generalize towards the same result. Furthermore, we observe that the correlation to the random classifier is lower for all models compared to the users (yellow bars below magenta). This demonstrates that model training reduces the noise in the classification.

### Prediction of foci sizes by our algorithm sizes is fairly close to some users but is of limited use depending on image resolution

SGs in our dataset are as little as 4-5px in radius. A deviation of 1px (or 40.7nm) therefore already constitutes a 25% deviation. (For data input the users worked on an upscaled image of the cell allowing for more control, but blurry images).

Under these considerations, the results of the size prediction are fairly close to the users with a tendency to overestimate the area. A further improvement is achieved by adjusting a parameter to increase or decrease all foci sizes by a parameter in the UI of our software **(Sup. Video 4)** - which in the best case corresponds to shifting the results in **Figure 5** upwards by their mean. Other solutions like a separate model for size prediction are feasible, however size of foci in absolute terms is often not informative by itself. Instead, detecting relative changes in foci size is more often informative, for which the problem of overestimating the size does not really matter.

**Figure 5:**
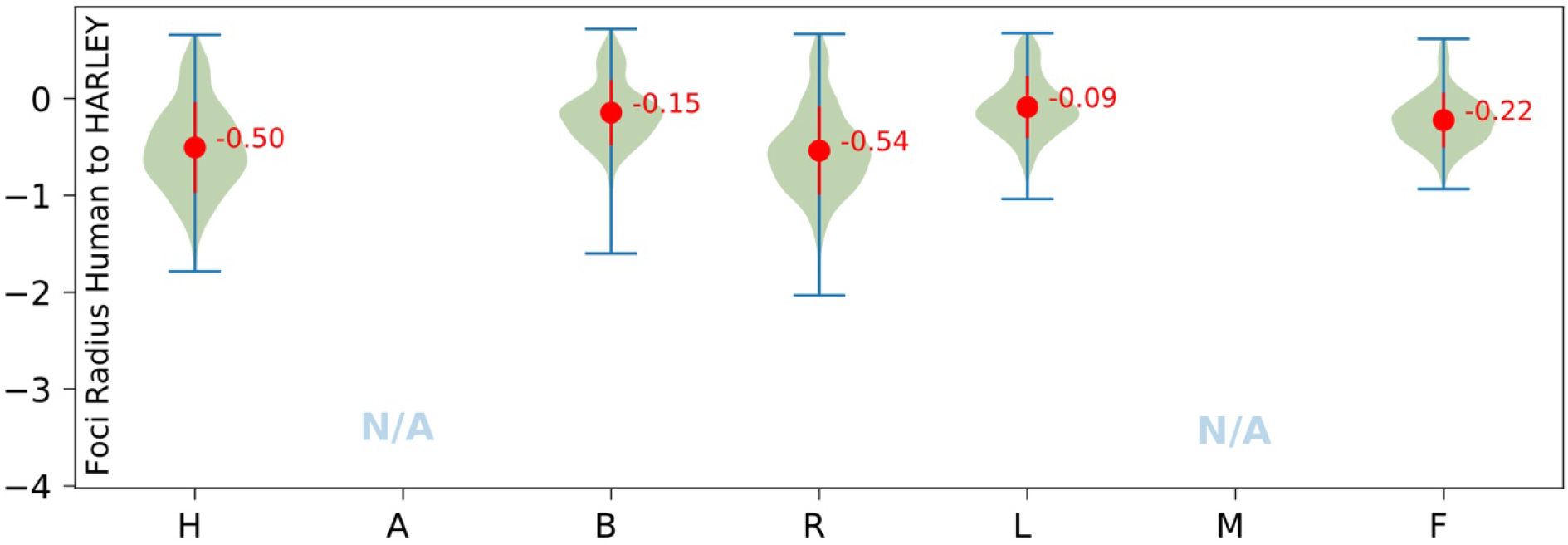
Foci radii determined by HARLEY typically exceed user-defined radii. 0 corresponds to equal sizes. Negative values correspond to an overestimation on the side of the model. Since foci are not round, radiae correspond to circles of same area as selected foci.

### A stress granule training set >15 cells achieves effective user SVM modelling

and we generated 30 samples for each subset size. The subset size is measured in number of foci candidates with the average cell containing 13.2 candidates.

While the performance of the model keeps increasing, a training set of 100-200 foci candidates (on average less than 15 cells in our standardized NaN3 dataset) is sufficient for the model to learn most users (**Figure 6**). In practice, labelling 15 cells takes not much more than 5-10 minutes with our software. Training times for the support vector machine itself for such small datasets are on the magnitude of milliseconds to seconds (for determining optimal hyper parameters), making this approach highly practical for users with relatively consistent scoring of foci.

**Figure 6:**
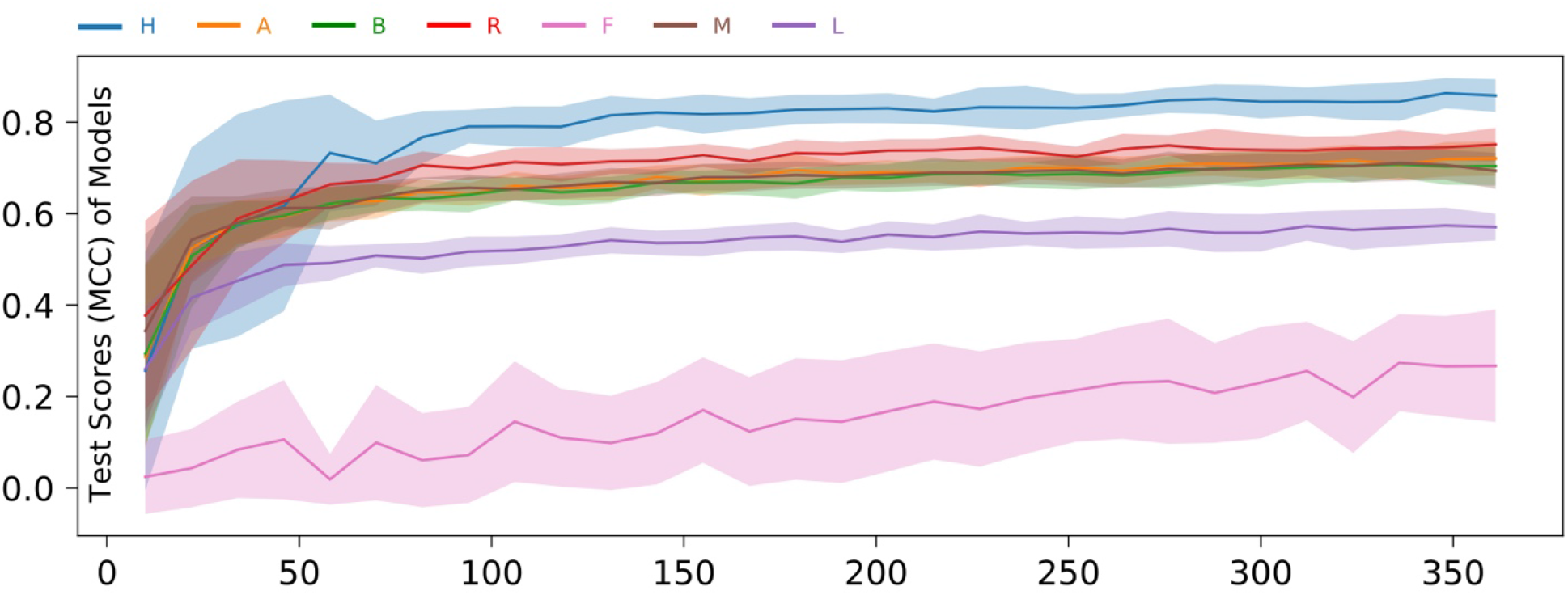
Model performance by size of dataset (in foci). Model hyper parameters are not optimised. Shaded areas show one standard deviation above and beyond the mean determined from 30 repeats for each setting.

## DISCUSSION

We have shown that user variability is a significant problem in scoring of somewhat ambiguous cellular foci such as SGs. In a group of just 7 users, variability approached 4-fold, with user reproducibility of foci identification only reaching a maximum of about 80%. This casts significant doubt on interpreting marginal differences in SG microscopy data generated by different labs, users and even within studies where only a single user has been responsible for quantification. To counteract such variability, and aid meaningful data comparisons, we developed HARLEY to facilitate rapid and consistent identification of yeast cell boundaries and foci using a novel contour-loop based approach. Our approach is much more invariant to foci shape (i.e., works equally well for round or elongated shapes) and deals with clumped foci in an elegant implicit fashion. We have shown that our data training approach can rapidly generate a model that will allow users to consistently and rapidly quantify foci of interest, including via a batch mode. The ability of users to reproduce their own results is indicative of model performance with the best results coming from users who can reproduce their own results with ~80% correlation, which is not surprising, since human quantification inconsistencies accumulate in the training dataset. This greatly facilitates data analysis for SGs and PBs that we have previous reported (Buchan et al., 2010).

Many popular software tools currently used for quantification of foci in microscopy data enable practical parameter-constrained solutions, that are very sensitive to a technical understanding of the interplay and influence of these parameters *(Schneider 2019)* or produce poor results on noisy data. A model training-based approach solves these issues by asking the user to define the desired outcome rather than parameters that lead to it. Biologists are often familiar with software like ImageJ, that while having a diverse array of uses, may become limited for high-throughput analysis of microscopy data, owing to a somewhat inflexible structure and the need for manual input.

The chief advantage of HARLEY is a clear, UI guided process that allows users with minimal technical expertise to quickly create consistent and reliable quantification results. The ability to share trained models allows reproduction and comparison of results reported by other labs, which we hope will increase the quality of reported data. While we are aware that deep learning approaches may generate better results, no deep learning approach can be trained in milliseconds using only a small number of cells. Thus, we anticipate that HARLEY will benefit the community and encourage more biologists to use automated software for their quantification.

To improve our approach for foci detection the feature space likely needs to be extended. A straightforward solution would be to use a feature extraction CNN to generate the feature vectors for the SVM that are most relevant to the user or generate extensive datasets using a variety of filters. While the number of weights in SVMs only grow with the dimension of the feature space (13 in our case and 2 hyper parameters), CNNs require learning of weights many orders of magnitudes above that (with values of 10^5^-10^6^ being not uncommon), which increases demands on time and computational power significantly. While a gain on accuracy is possible, the quality of learning data may be more problematic due to strong noise in human labeling. A reasonable trade off on accuracy and practical usability must be found with the application of these approaches. A potential gain in consistency might be achieved by using quantifications from multiple human labelings combined into one model, to assess a common denominator of what humans deem foci.

More work should be conducted to assess how well a trained model solves the task of foci detection. However, it should be stressed that comparing a model to a user baseline is somewhat limited, due to inconsistencies across human users. As in our case, some humans can be “learned” very well (H), while others are very hard to learn (F). The user “baseline” is therefore a spectrum which is not surprising, since there is no clear definition of what foci are. One solution is the use of standardized datasets and performance measures that the scientific community generally lacks. We are aiming at developing a more extensive dataset for SGs and P-bodies, as well as other cellular structures amenable to contour-based identification, in the future. The current dataset is freely available already at https://github.com/lnilya/harley.

While our model readily learns some users the performance for others is poor, which arguably could be due to the noisiness of their classification. If a user can reproduce their own result only to a 60-80% correlation arguably any generalizing model will not learn to more than this precision, making it indicative of an upper bound. While we did not have enough data to fully cement this notion, the results strongly suggest that users should assess their own performance to determine the quality of their quantification reproducibility, and thus how well their trained model is expected to perform. In future updates of HARLEY, we plan to incorporate this feature.

Our initial studies focused on SGs, since they are a challenging organelle to quantify with a significant time burden. We anticipate however that HARLEY will be readily transferable to the quantification of other organelles whose foci are approximately convex in nature and do not overlap (e.g., P-bodies, kinetochores, microtubules) Adjustments will have to be made for morphologically more complex shapes such as mitochondria. As HARLEY is developed further, other organelles are likely to be included in the future. Our goals for HARLEY are to develop it into an intuitive platform for biologists working with microscopy data. We also plan to integrate features like colocalization and allow users to assess variability to others and themselves.

Finally, since HARLEY is based on a problem agnostic pipeline-based framework that we are developing separately, interested researchers are encouraged to reach out and participate in the development of a new platform for delivering simple, intuitive UIs to any algorithms that would be too complex or unwieldy as a plugin to existing software. Since HARLEY is based on ReactJS, the usability, ease of development and cross platform compatibility are among its greatest strengths. We hope to encourage the community to build or retrofit their algorithms into beautiful easy to understand tools using this framework, benefiting the scientific community.

